# Estimation of cellularity in tumours treated with Neoadjuvant therapy: A comparison of Machine Learning algorithms

**DOI:** 10.1101/2020.04.09.034348

**Authors:** Mauricio Alberto Ortega-Ruíz, Cefa Karabağ, Victor García Garduño, Constantino Carlos Reyes-Aldasoro

## Abstract

This paper describes a method for residual tumour cellularity (TC) estimation in Neoadjuvant treatment (NAT) of advanced breast cancer. This is determined manually by visual inspection by a radiologist, then an automated computation will contribute to reduce time workload and increase precision and accuracy. TC is estimated as the ratio of tumour area by total image area estimated after the NAT. The method proposed computes TC by using machine learning techniques trained with information on morphological parameters of segmented nuclei in order to classify regions of the image as tumour or normal. The data is provided by the 2019 SPIE Breast challenge, which was proposed to develop automated TC computation algorithms. Three algorithms were implemented: Support Vector Machines, Nearest K-means and Adaptive Boosting (AdaBoost) decision trees. Performance based on accuracy is compared and evaluated and the best result was obtained with Support Vector Machines. Results obtained by the methods implemented were submitted during ongoing challenge with a maximum of 0.76 of prediction probability of success.

## 1 Introduction

Breast cancer NAT therapy has been used as a locally treatment for breast-conserving surgery [1], it provides prognostic and survival information [2] and is also used to determine a rate of local recurrence [3]. Efficacy of NAT is determined by means of the pathological complete response (pCR) but an accurate assessment of pCR is needed. It has been proposed [4] the Residual Cancer Burden (RCB) as a long term prognosis method. RCB is supported by two metrics: residual Tumour Cellularity (TC) within the Tumour Bed (TB) and assessment of lynph nodes. RCB is scored in a continuous value but is further categorised in four classes RCB-0 to RCB-III. Then TC is a key parameter for RCB computation. Currently, TC is assessed by an eye-balling routine estimating the proportion of TB and is compared to a standard sketch reference. Values are assigned manually and are rounded to the nearest tenth percent value. This procedure is time-consuming and requires an experienced well trained pathologist. With the recent development of computational methods based on the use of artificial intelligence and computer vision algorithms combined with the use of whole slides scanners, Digital Pathology can provide a more efficient and accurate method to address this problem, as it has been proved in computer aid diagnosis (CAD) [5], as in other clinical application tasks like segmentation of a region of interest (ROI), mitosis detection, gland segmentation, and even new clinic pathological relationships with a specific image morphological behaviour [6].

Machine learning algorithms [7] are based on the use of extracted image features to train the method with known data, by means of digital pathology steps: color separation, nuclei segmentation and morphological features extraction [8]. However, Deep learning methods based on Convolutional Neural Networks (CNNs) [9] have the advantage of train the net directly with the images without feature extraction. TC estimation problem was addressed by Peikari [10] who proposed a method based on nuclei segmentation and morphological parameters extraction used to train machine learning algorithms to classify cells as benign or malign. Akbar [1] compares this technique with deep learning and evaluates performance and also Pei [11] implemented a direct method based on transfer learning approach. Also during the 2019 SPIE Breast Path Q Challenge called for development of automated TC algorithms.

The present paper describes a method for an automated TC estimation based on machine learning methods using key selected best correlated parameters to TC. A prior correlation analysis of morphological features at nuclei, regional and global image level were done and only highest correlated parameters were employed to train the method.

## 2 Materials and Methods

### 2.1 Materials

The data set for this study comes from the Breast SPIE Challenge 2019 collected at the Sunnybrook Health Sciences Centre, Toronto. It comprises a set of 67 whole slide image (WSI) from post-NAT patients of Breast Cancer stained with Hematoxilyn and Eosin (H&E) [10]. The specimens were handled according to routine clinical protocols and WSIs scanning was performed at 20*×* magnification (0.5*µ* m/pixel). There is a training set of 2579 patches of size 255 *×* 255 extracted from the above WSIs which has a reference standard from 2 pathologists who classified cellularity on a score scale from 0 to 1. This manual classification is used as the reference Ground Truth (GT) of this study. Fig. 1 shows samples of these images with TC values from 0 to 1.0. At each sample there is a caption information with pathologist classification and results obtained by the methods. Also there is a set set of 1119 images provided to analyse and submit results to the contest, with no information on TC. A third set of selected images was provided by the challenge committee with pathology annotations indicating: normal, malignant and lymphocytes. All these three data sets were used in this work.

**Fig. 1.**
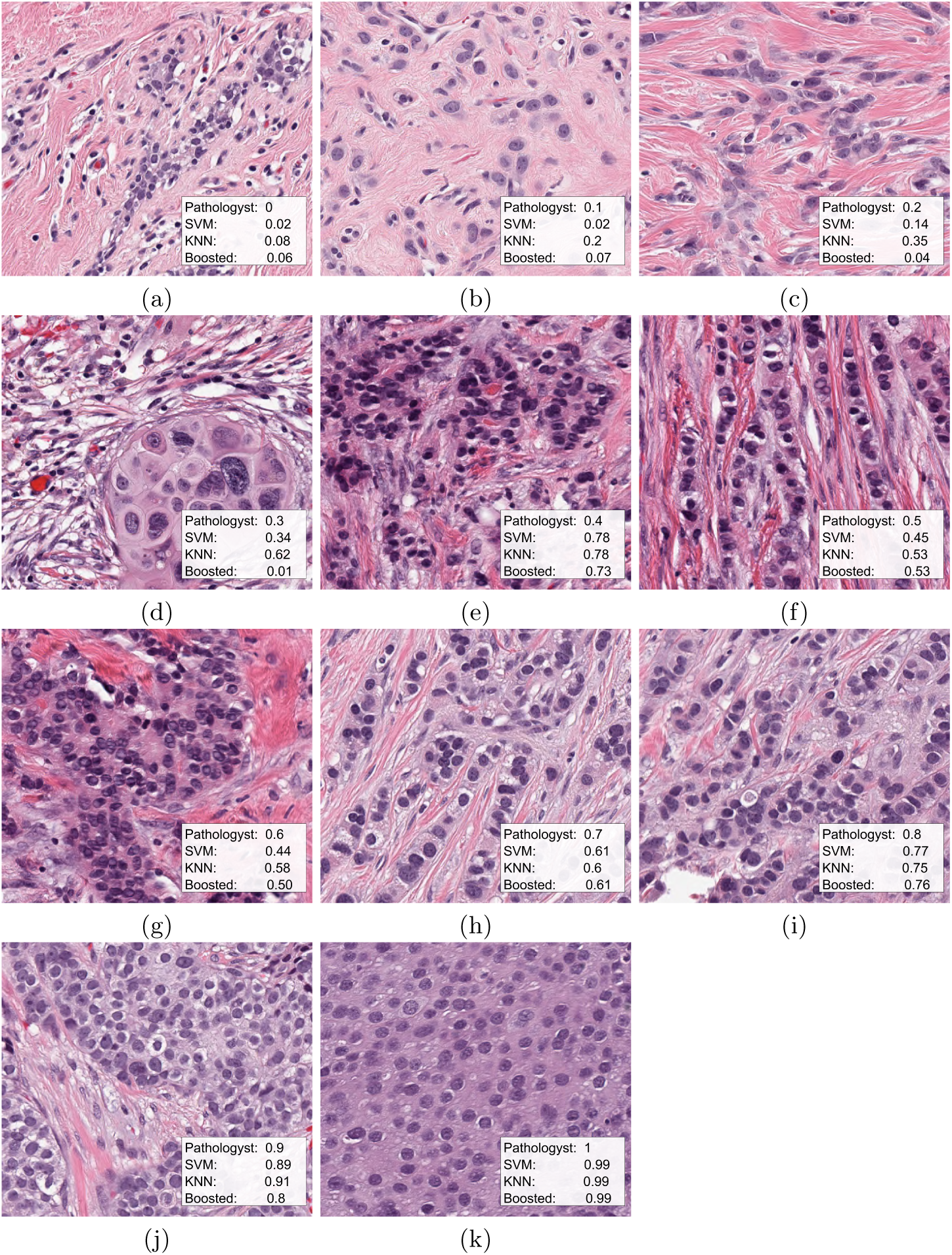
Several representative samples of breast cancer NAT tissue stained with Hematoxilyn and Eosin (H&E). The images correspond to different values of TC, from 0 to 1. Each image contains four annotations; a classification as assess by a pathologist and the results by each of the methods implemented in this paper.

### 2.2 Methods

The method proposed in this study implements traditional digital pathology steps [8]: color separation, nuclei segmentation, binarisation and feature extraction from nuclei, regional and global areas of the image. This method uses the same methodology as described by Peikari hand engineering processing but with the advantage of being trained only by key selected morphological image features that are strongly correlated with TC.

A special image processing software was developed to analyse a large amount of pathology images. A master control routine selects and processes one by one the corresponding image and also saves the result in an output data file. A supervised operation mode of the software is used to train machine learning algorithms to determine a predicting function for classifying cells into the two corresponding categories: benign or malign. The unsupervised mode process the full image set and after extraction of image features it also executes the corresponding predicting function that classifies every segmented cell. Finally, TB region is estimated in order to compute TC as a unique value for the corresponding patch image.

The full method software was implemented in Matlab ® (The Mathworks™, Natick, USA) 2019b version, using digital image processing, statistical and machine learning toolboxes.

The method process is depicted in Fig. 2. First, nuclei is segmented and its corresponding key parameters are extracted (Fig. 2a). After the prediction function is computed nuclei is classified in either benign (green) or malign (red) (Fig. 2b). Next, an estimation of of full cell cytoplasm by morphological dilation is shown as the white circles around malign cells (Fig. 2c). The full cellularity detected region is shown in Fig. 2d. Finally TC is computed by the ratio of cellularity area by global image area. A continuous TC value is obtained by the method in contrast with the selected TC by pathologist in tenths intervals, a continuous value can offer a better accuracy, which indicate TC should be reformulated.

**Fig. 2.**
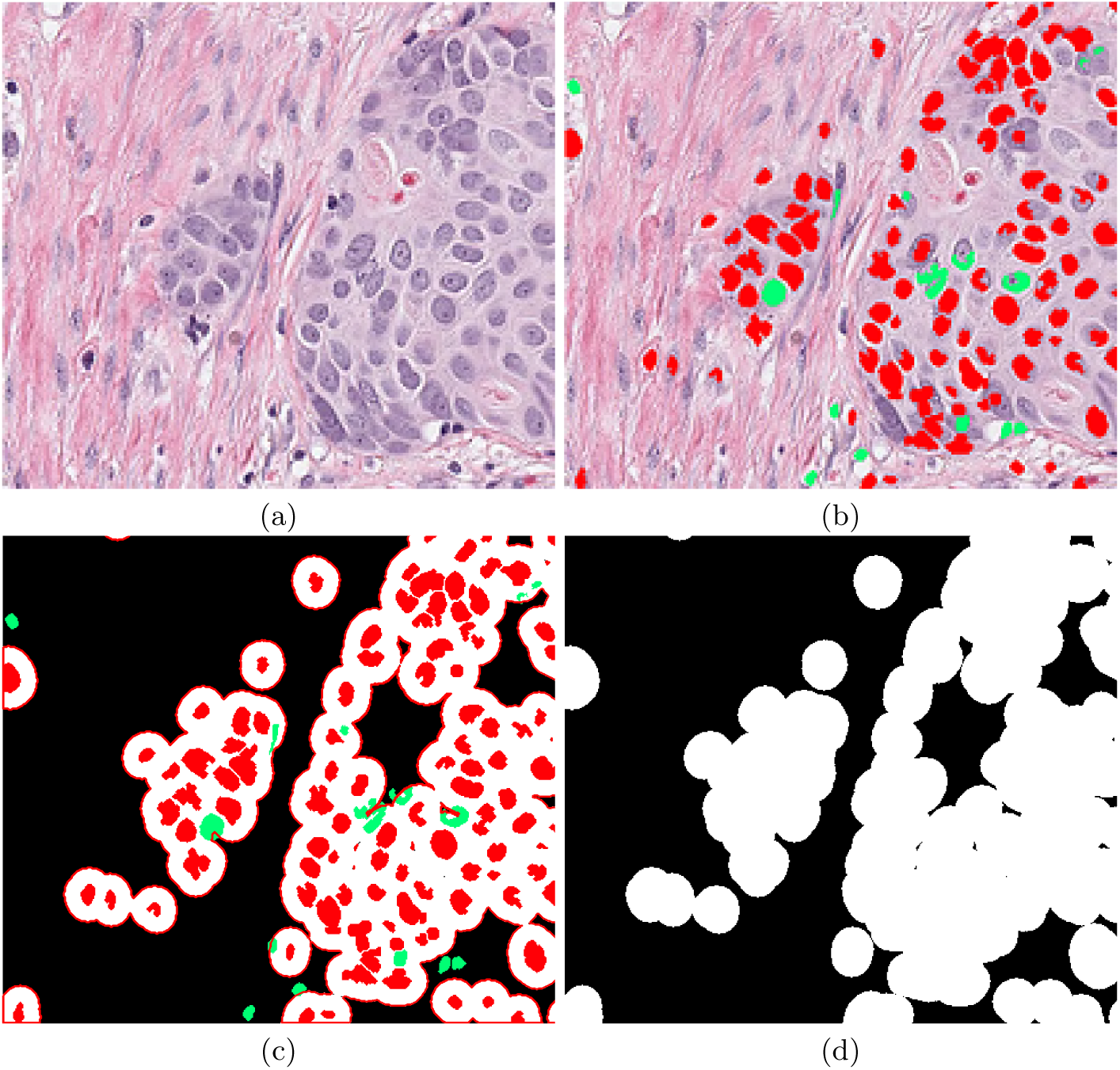
Visual description of the method. (a) The original image, the image is segmented and key parameters are computed, then a classification predictor estimates either malignant or benign cells, shown in red and green respectively in (b). A dilation of segmented malignant nuclei estimates full cytoplasm of every detected malignant cell (c) and overall cellularity region is shown in white in (d).

#### Data pre-processing

First, a correlation analysis of main morphological parameters with residual TC was obtained. Results indicated 22 parameters have a strong correlation with cancer cellularity and only these key parameters were used to train machine learning methods. These 22 parameters were obtained from the nuclei morphological analysis, from a regional analysis of the nuclei surrounding neighborhood and from the full image. The relevance of this paper is a machine learning training procedure with only key data for a better performance of predicting function. Three algorithms were employed to classify cells as benign or malign: Support Vector Machines, Nearest K-Network and Ad-aBoost. These algorithms were trained during a pre-processing step to generate a predictive function to classify each cell.

#### Training of machine learning algorithms

Support Vector Machines (SVM) were presented as a training algorithm for optimal margin classifier [12], and it is based on determination of a decision function of pattern vectors of *x* of dimension *n* classifying in either A or B, which means benign or malign. The input is a set of *p* examples of *x*_*i*_, in this case the 22 strongest correlated features extracted. Relationship of the strongest related parameters with TC are presented in Fig. 3, for the optimal cases a clear separation can be observed in figures.

**Fig. 3.**
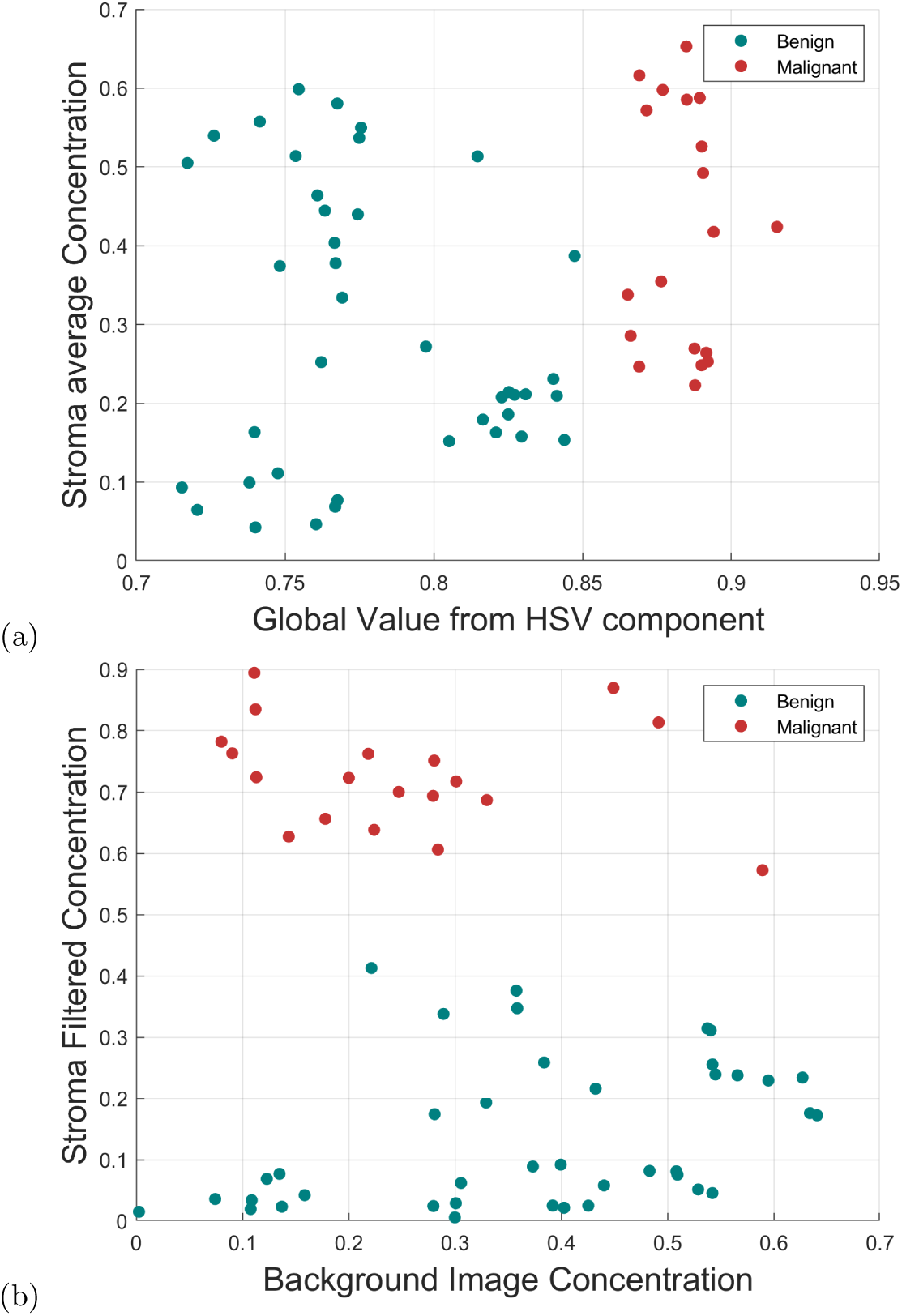
Morphological relationship between strongest correlated parameters. The 4 strongest morphological correlated parameters (from the 22 detected) with TC are compared, which are: 1) an average stroma concentration determined by a pink colour filter, 2) average Value V from HSV global image, 3) stroma concentration from pink colour separation image as well as 4) background colour concentration from the image. Image indicates key parameters have a clear classification category

Then a decision function is expressed as *D*(*x*) = Σ*w*_*i*_*ϕ*_*i*_(*x*)+*b* in which *ϕ*_*i*_(*x*) is a predefined function of *x* and *w*_*i*_ and *b* are adjustable parameters. Assuming that there is a separation margin *M* between both classes the algorithm determines vector *w* that maximizes M attained to minimise *y*_*k*_*D*(*x*_*k*_) = *M*\*. This algorithm was selected as it has been successfully reported in different pathology studies included TC estimation [11].

K- Nearest Neighbour method [13] for machine learning classification was also selected because is one of the most popular algorithms within clustering and data classification methods, in this case, grouping between benign and malign classes. This method is based on the following: let assume *T* is a two dimensional set of vectors with their corresponding elements *{x*_*i*_, *y*_*i*_ *}*, which is used as the training set, then a new sample, say *x* = *un* is given, and the algorithm has to estimate the class that this sample belongs. The algorithm starts from the simplest case *K* = 1 the sample closest to *u* is found and set *v* = *y* where y is the closest class of the nearest neighbour sample. To extend this idea to the *K* − *NN* dimension, the nearest *K* neighbours elements need to be found and a majority decision rule classifies the new sample. Distance is measured by the euclidean distance.

AdaBoost is an algorithm that has a best performance on binary classification problems, for that reason it was selected to classify between malign and benign nuclei cell. AdaBoost is a decision tree type learning algorithm [14], which is a predictive model that starts from observations of a certain item represented by branches and goes to conclusions about item target value (leaves). The algorithm is as follows. It starts assuming same weight to each training point 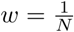. Next error rate for weak classifier is calculated as *e* = Σ*w*_*i*_ and those classifiers with lowest error rate are picked up and it computes a voting power 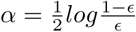. The classifier is appended in the assemble and decides if classifier is good enough. Weights of previous wrong classifiers are updated by new *w* as follows 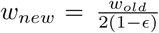 for point classified correctly and 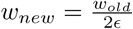 for incorrectly case.

## 3 Results

An automated estimation of TC was computed from two test data sets. Three prediction functions trained by machine learning algorithms were determined to be used with the automated processing software of breast cancer images that classifies cells and computes TC. The method was tested with a training set of 2579 images already classified by a pathologist with a TC value. Also it was tested with the 1119 images for submission of SPEI Breast Challenge, with an unknown TC value. Fig. 4 shows the statistical behaviour of the method’s result for the training set as scatter plots and Fig. 5 the same results as boxplots. Fig. 4(a) is the corresponding dispersion plot for SVM prediction function in which every black circle represents a single patch image computation result and the line indicate the true value. The red spot is the median point. Fig. 4(b) shows the results for KNN algorithm and Fig. 4(c) for AdaBoost. Initially, to evaluate the performance mean square error of the three results were computed and presented in first row of Table 1. Dispersion plot indicates the method approximates to the pathologist classification assignment. Results have a better approximation at higher cellularity values (*TC >* 0.70) and performs well with KNN algorithm. Also around the middle region (0.4 *< TC <* 0.6) has a good approximation with AdaBoost. At low cellularity values (*TC <* 0.3) three methods present deviation, with its higher at cellularity zero, which correspond to images with only benign nuclei cells. According to Minimum Square Error (MSE) SVM performs better overall the cellularity region. This result can be validated by a visual inspection of boxplots of Fig. 5 in the three cases there is a positive correlation between the actual celullarity (horizontal) and the estimated cellularity (vertical). However, SVM shows less dispersion, especially in the lower values of cellularity as compared with the other two techniques.

**Table 1.** Performance evaluation of the method. Mean square error (MSE) was computed of the result obtained over the full train data set, using pathologist’s as a ground truth (GT) reference. Prediction probability determined after data submission at the contest. Confidence intervals are presented in brackets.

| Performance Metric | SVM | KNN | AdaBoost | Data Set |
| --- | --- | --- | --- | --- |
| MSE | 0.0352 | 0.0655 | 0.08 | Training Data |
| Prediction Probability | 0.766 [0.72,0.81] | 0.754[0.70, 0.80] | 0.712[0.67, 0.75] | Test Data |

**Fig. 4.**
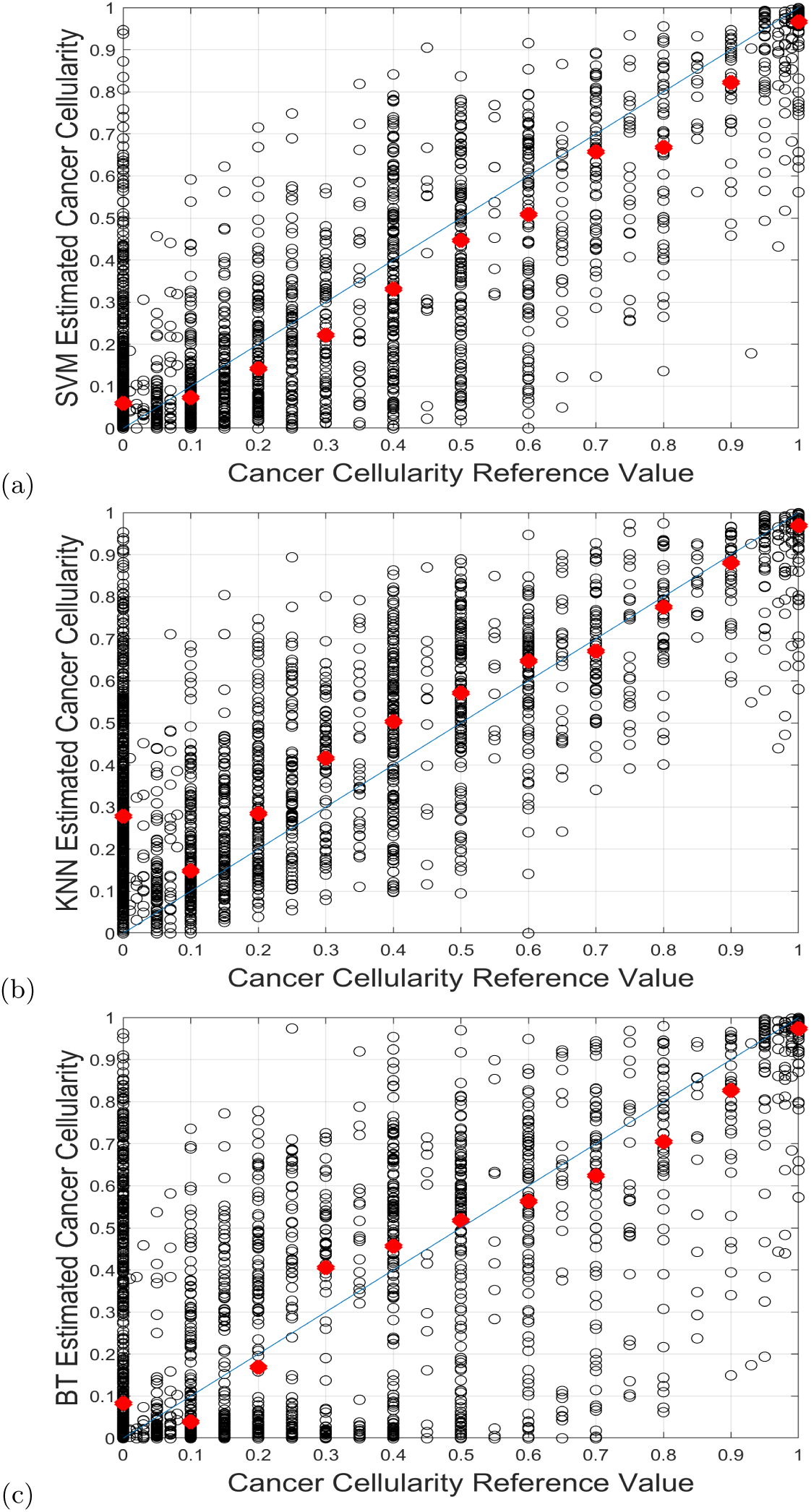
Results of implementation on Training Data shown as scatter plots over the training set of 2579 patches. Horizontal axis is the reference cellularity value and vertical axis is the value computed by the method. The line represents the true value, every single circle is an image patch result and the red spot is the median of a single Cellularity value.

**Fig. 5.**
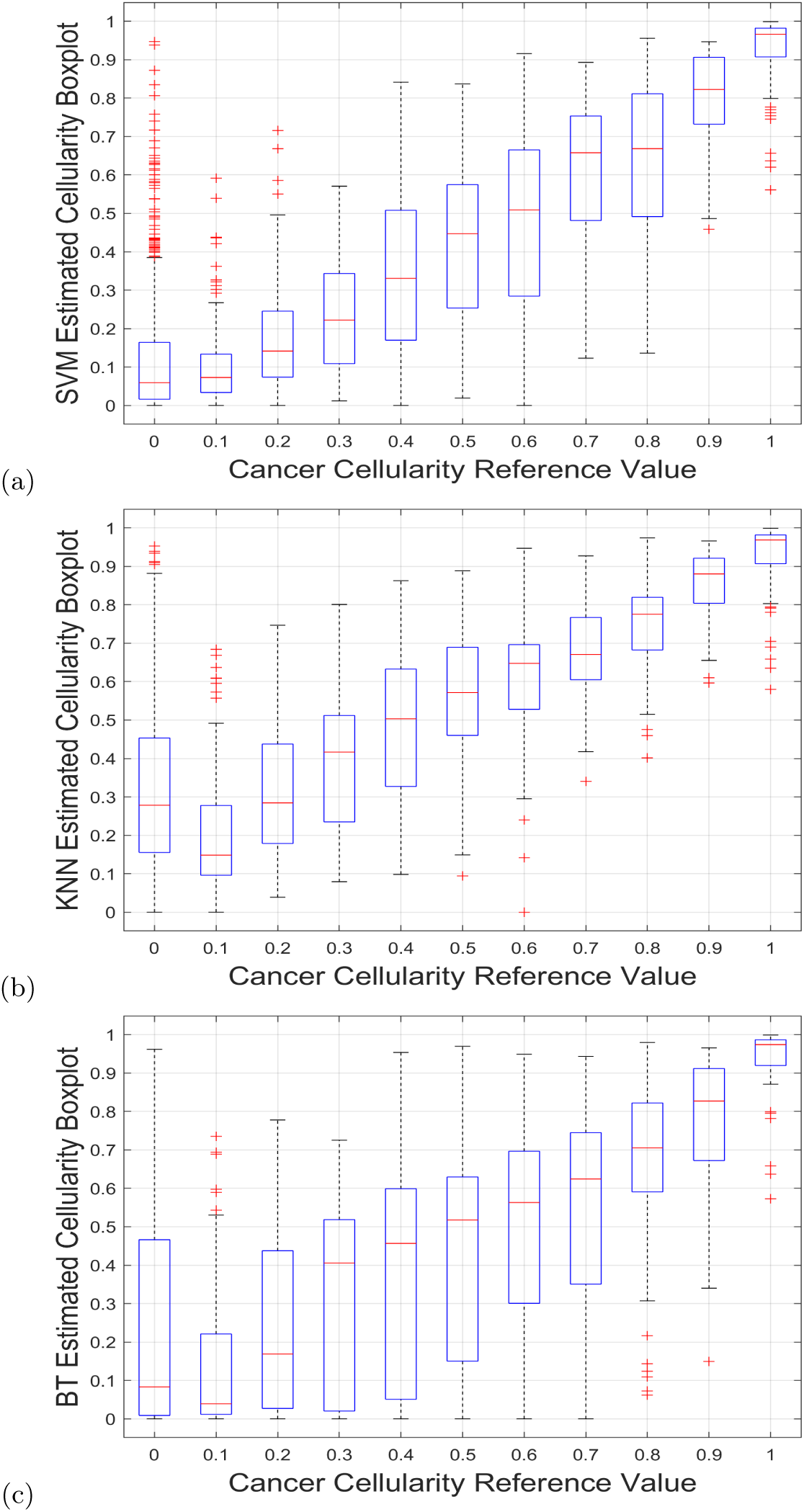
Results of implementation on Training Data shown as boxplots. Every boxplot corresponds to the statistical result of all the same TC value results. Large boxplot indicate large deviation. A visual inspection indicates the lowest deviation corresponds to SVM method in (a).

Finally, the method results from contest challenge data were evaluated by means of the prediction probability results given by the challenge and are presented at second row of Table 1 with a maximum of *P*_*k*_ = 0.76 obtained with SVM algorithm.

## 4 Discussion

An automated processing software for TC computation is presented in this paper. The methodology was tested to process automatically breast cancer images under Neoadjuvant treatment. To determine tumour regions, three different machine learning algorithms for cell classification were evaluated and compared. The **best result was obtained with SVM algorithm**. The relevance of this paper is a selection of a key parameters to train the algorithms which results in a better performance of similar techniques, however reported deep learning algorithms outperforms this result, which is a motivation to explore these technique in the future.

